# Mechanistic insights into the function of 14-3-3 proteins as negative regulators of brassinosteroid signaling in Arabidopsis

**DOI:** 10.1101/2023.10.13.562204

**Authors:** Elsa Obergfell, Ulrich Hohmann, Andrea Moretti, Michael Hothorn

## Abstract

Brassinosteroids (BRs) are vital plant steroid hormones sensed at the cell surface by a membrane signaling complex comprising the receptor kinase BRI1 and a SERK-family co-receptor kinase. Activation of this complex lead to dissociation of the inhibitor protein BKI1 from the receptor and to differential phosphorylation of BZR1/BES1 transcription factors by the glycogen synthase kinase 3 protein BIN2. Many phosphoproteins of the BR signaling pathway, including BRI1, SERKs, BKI1 and BZR1/BES1 can associate with 14-3-3 proteins. In this study, we use quantitative ligand binding assays to define the minimal 14-3-3 binding sites in the N-terminal lobe of the BRI1 kinase domain, in BKI1, and in BZR1 from *Arabidopsis thaliana*. All three motifs require to be phosphorylated to specifically bind 14-3-3s with mid- to low micromolar affinity. BR signaling components display minimal isoform preference within the 14-3-3 non-ε subgroup. 14-3-3ƛ and 14-3-3ω isoform complex crystal structures reveal that BKI1 and BZR1 bind as canonical type II 14-3-3 linear motifs. Disruption of key amino acids in the phosphopeptide binding site through mutation impairs the interaction of 14-3-3ƛ with all three linear motifs. Notably, quadruple loss-of-function mutants from the non-ε group exhibit gain-of-function brassinosteroid signaling phenotypes, suggesting a role for 14-3-3 proteins as overall negative regulators of the BR pathway. Collectively, our work provides further mechanistic and genetic evidence for the regulatory role of 14-3-3 proteins at various stages of the brassinosteroid signaling cascade.

## Introduction

Brassinosteroids are a class of polyhydroxylated plant steroid hormones (Grove et al., 1979) perceived at the plasma membrane by the leucine-rich repeat membrane receptor kinase BRASSINOSTEROID INSENSITIVE 1 (BRI1) (Clouse et al., 1996; Li and Chory, 1997; He et al., 2000; Wang et al., 2001; Hothorn et al., 2011; She et al., 2011; Hohmann et al., 2018b). Brassinosteroid binding to BRI1 enables binding of leucine-rich repeat co-receptor kinases of the BRI1 ASSOCIATED KINASE 1 / SOMATIC EMBRYOGENESIS RECEPTOR KINASE (BAK1/SERK) family (Schmidt et al., 1997; Li et al., 2002; Nam and Li, 2002; Gou et al., 2012; Santiago et al., 2013; Hohmann et al., 2018b). In the absence of BRs, SERKs can constitutively bind to BAK1-INTERACTING RECEPTOR-LIKE KINASE (BIR) receptor pseudokinases, negative regulators of BR signaling (Imkampe et al., 2017; Hohmann et al., 2018a). Brassinosteroid-induced heterodimerisation of BRI1 with a SERK enables trans-phosphorylation of their cytoplasmic dual-specificity kinase domains (Friedrichsen et al., 2000; Oh et al., 2000, 2009; Bücherl et al., 2013; Wang et al., 2008; Bojar et al., 2014). BRI1 kinase activation leads to the phosphorylation and dissociation of the largely unstructured inhibitor protein BRI1 KINASE INHIBITOR 1 (BKI1) from the BRI1 kinase domain, accompanied by the relocalisation of BKI1 from the plasma membrane into the cytosol (Wang and Chory, 2006; Jiang et al., 2015; Wang et al., 2011, 2014; Jaillais et al., 2011; Novikova et al., 2022). Cytoplasmic BRI1 signaling results in inactivation of the glycogen synthase kinase 3 family protein BR INSENSITIVE 2 (BIN2) (Li et al., 2001; Li and Nam, 2002; De Rybel et al., 2009; Kim et al., 2009) by the protein phosphatase BRI1 SUPPRESSOR 1 (BSU1) (Mora-García et al., 2004; Kim et al., 2009). Downstream of BIN2, reduced phosphorylation of the transcription factors BRASSINAZOLE-RESISTANT 1 (BZR1) / BRI1-EMS-SUPPRESSOR1 (BES1) promotes their nuclear localisation and mediates BR-responsive gene expression (He et al., 2002; Yin et al., 2002; Zhao et al., 2002; Vert and Chory, 2006; Ryu et al., 2007, 2010; Tang et al., 2011; Nosaki et al., 2018).

Different phosphoproteins involved in BR signaling have been previously reported to bind 14-3-3 proteins, a family of dimeric scaffolding proteins engaging in phosphorylation-dependent protein – protein interactions (Fu et al., 2000). 14-3-3s provide a conserved cup-shaped binding groove for phosphorylated protein ligands (Ballone et al., 2018). In plants, 14-3-3 proteins have been implicated in the regulation of cellular metabolism, ion and nutrient homeostasis (Ottmann et al., 2007; Denison et al., 2011; Xu et al., 2012; Yang et al., 2019; Gao et al., 2021a, 2021b; Jiang et al., 2023), in plant immunity (Konagaya et al., 2004; Chang et al., 2009; Stanislas et al., 2009), and in different light and hormone signaling pathways (Sullivan et al., 2009; Sirichandra et al., 2010; Taoka et al., 2011; Tseng et al., 2012; Gao et al., 2014; Huang et al., 2018; Prado et al., 2019; Reuter et al., 2021; Waksman et al., 2023). 14-3-3 proteins have been previously reported to directly or indirectly associate with different BR signaling components including BRI1 and SERKs (Rienties et al., 2005; Karlova et al., 2006; Chang et al., 2009), BKI1 (Wang et al., 2011), BSU1 (Chang et al., 2009), BIN2 (Kim et al., 2023) and BZR1/BES1 (Gampala et al., 2007; Bai et al., 2007; Ryu et al., 2007, 2010; Wang et al., 2013). Here we report a quantitative biochemical approach to define and fine map 14-3-3 interaction sites for different BR pathway components, and a reverse genetic analysis of the contribution of 14-3-3 isoforms to BR signaling.

## Results

### BRI1, BKI1 and BZR1 contain linear 14-3-3 binding motifs

We recombinantly expressed and purified Arabidopsis 14-3-3 isoform kappa (14-3-3κ) and tested for interaction with the globular domains of different BR signaling components by isothermal titration calorimetry (ITC). As 14-3-3 proteins selectively bind phosphorylated substrates, we auto- and transphosphorylated BRI1 and BAK1 cytoplasmic domains as well as BSK1 (Fig. 1A), as previously described (Oh et al., 2000; Kim et al., 2009; Bojar et al., 2014; Wang et al., 2014). No specific binding was detected for the isolated BRI1^814-1196^ cytoplasmic domain (residues 814-1196) after incubation with the BAK1^250-615^/SERK3 cytoplasmic domain (residues 250-615), for BAK1^250-615^ after incubation with BRI1^814-1196^, for BSK1 (residues 55-512) after incubation with BRI1^814-1196^ (Fig. 1A), for full-length BSU1 expressed in baculovirus-infected insect cells, or for BIN2 (residues 7-380) expressed either in *E. coli* or in insect cells (see Methods, Fig. 1A,D).

**Fig. 1.**
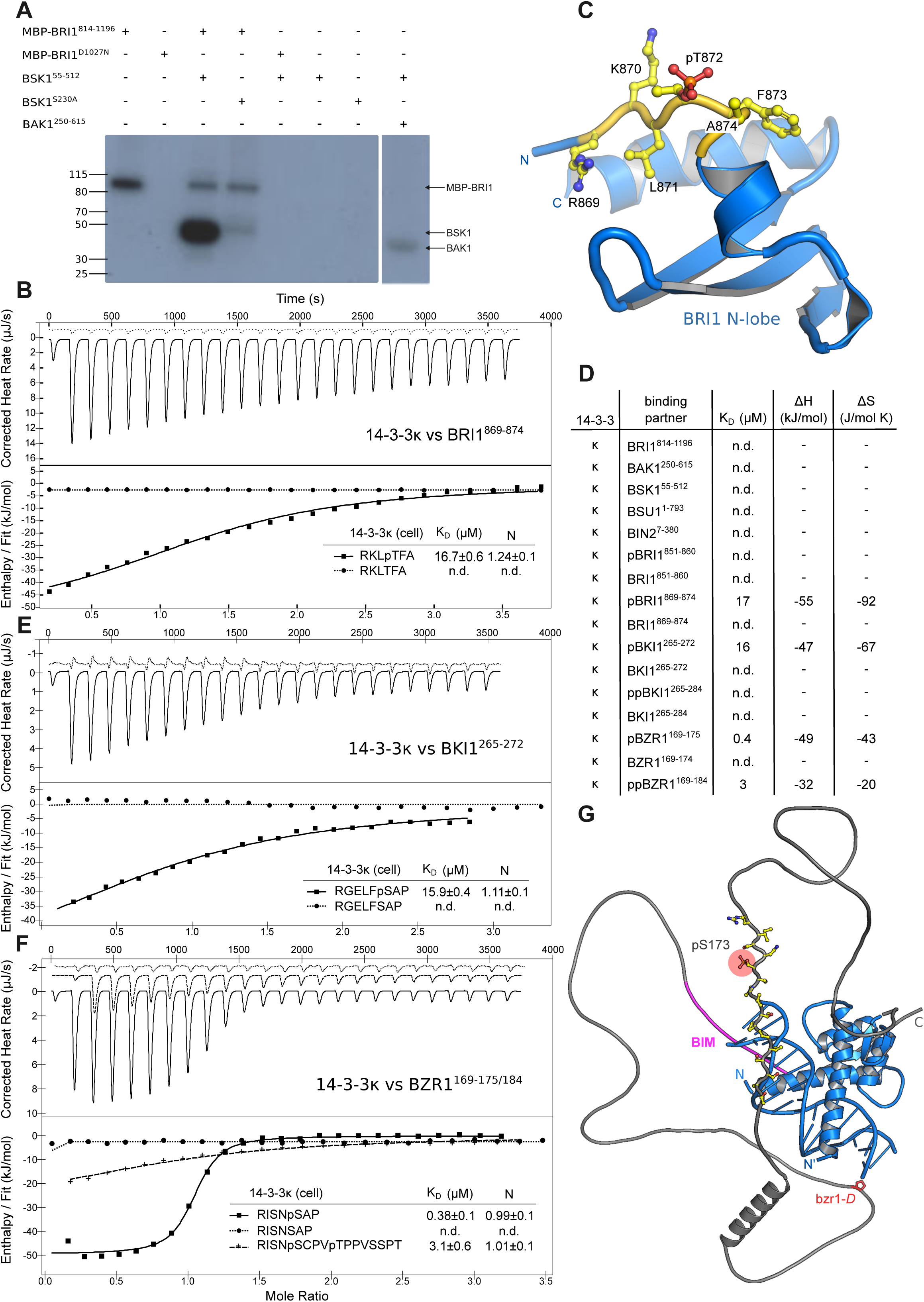
Short linear motifs in BRI1, BKI1 and BZR1 represent 14-3-3 binding sites. (A) *In vitro* transphosphorylation assay of BRI1 vs. BSK1. A Maltose-binding protein (MBP) fusion protein of wild-type AtBRI1 cytoplasmic domain (residues 814-1196; lane 1) but not the kinase inactive (Asp1027→ Asn) mutant version (lane 2), can efficiently trans-phosphorylate an AtBSK1 fragment covering residues 55-512 (lane 3). BSK1 phosphorylation by BRI1 mainly involves BSK1 Ser230 (lane 4), as previously shown (Tang et al., 2011). The sequence-related co-receptor kinase BAK1 (residues 250-615) is unable to phosphorylate BSK1 (lane 8). (B) Isothermal titration calorimetry of 14-3-3κ vs. a phosphorylated short linear motif located in the N-lobe of the BRI1 kinase domain (pBRI1^869-874^, continuous line). Shown are integrated heat peaks (upper panel) vs. time and fitted binding isotherms vs. molar ratio of peptide ligand (lower panel). No binding was detected for the unphosphorylated peptide (dotted line; n.d. no detectable binding). Table summaries for dissociation constants (K_D_) and binding stoichiometries (N) are shown (±fitting error). (C) Location of the 14-3-3 binding motif in the BRI1 kinase domain structure. Shown is a ribbon diagram of the N-lobe of the cytoplasmic kinase domain of AtBRI1 (in blue, PDB-ID 5lpb, residues 869-934) (Bojar et al., 2014), and the identified 14-3-3 binding motif (in yellow, residues 869-874) harboring phosphothreonine 872 (in bonds representation). (D) Table summaries of all ITC experiments performed with the 14-3-3κ isoform. Shown are dissociation constants (K_D_), binding enthalpy (ΔH) and entropy (ΔS). All binding stoichiometries were 2:2 with N∼1. Experiments were repeated at least twice. (E) ITC experiments performed for 14-3-3κ vs. the short linear motif in AtBKI1 (residues 265-272), plotted as in (B). (F) ITC experiments for 14-3-3κ vs. the short (residues 169-175, continuous line) and extended (residues 169-184, dashed line) linear motifs in AtBZR1. No binding was observed for the unphosphorylated peptide (dotted line). (G) Relative positions of the 14-3-3 binding site (in yellow), the BIN2 interacting motif (in magenta) and the bzr1-D missense mutation (Pro234→Leu) (Wang et al., 2002) mapped onto a AtBZR1 AlphaFold (Jumper et al., 2021) model (https://search.foldseek.com with ID Q8S307). The experimental BZR1 DNA binding motif – DNA complex structure (PDB-ID 5zd4) (Nosaki et al., 2018) is shown as a structural superposition (blue ribbon diagram).

Next, we used the eukaryotic linear motif (ELM) resource server (Puntervoll et al., 2003) to identify putative linear 14-3-3 binding motifs in BR cytoplasmic signaling components. Two such motifs were found in the juxtamembrane region of BRI1: Ser858 in KEALSIN (BRI1^851-860^) and Thr872 in the RKL<u>T</u>FA (BRI1^869-874^), both of which represent genuine BRI1 autophosphorylation sites (Oh et al., 2000; Wang et al., 2005). 14-3-3κ specifically bound the phosphorylated pBRI1^869-874^ but not the BRI1^851-860^ motif with mid micromolar affinity and with 2:2 (N=1, see below) binding stoichiometry (Fig. 1B,D, Table 1). This motif is located upstream of the catalytic BRI1 kinase core, where it folds into an additional β-strand that packs against the N-lobe of the kinase domain (Fig. 1C) (Bojar et al., 2014).

**Table 1.**
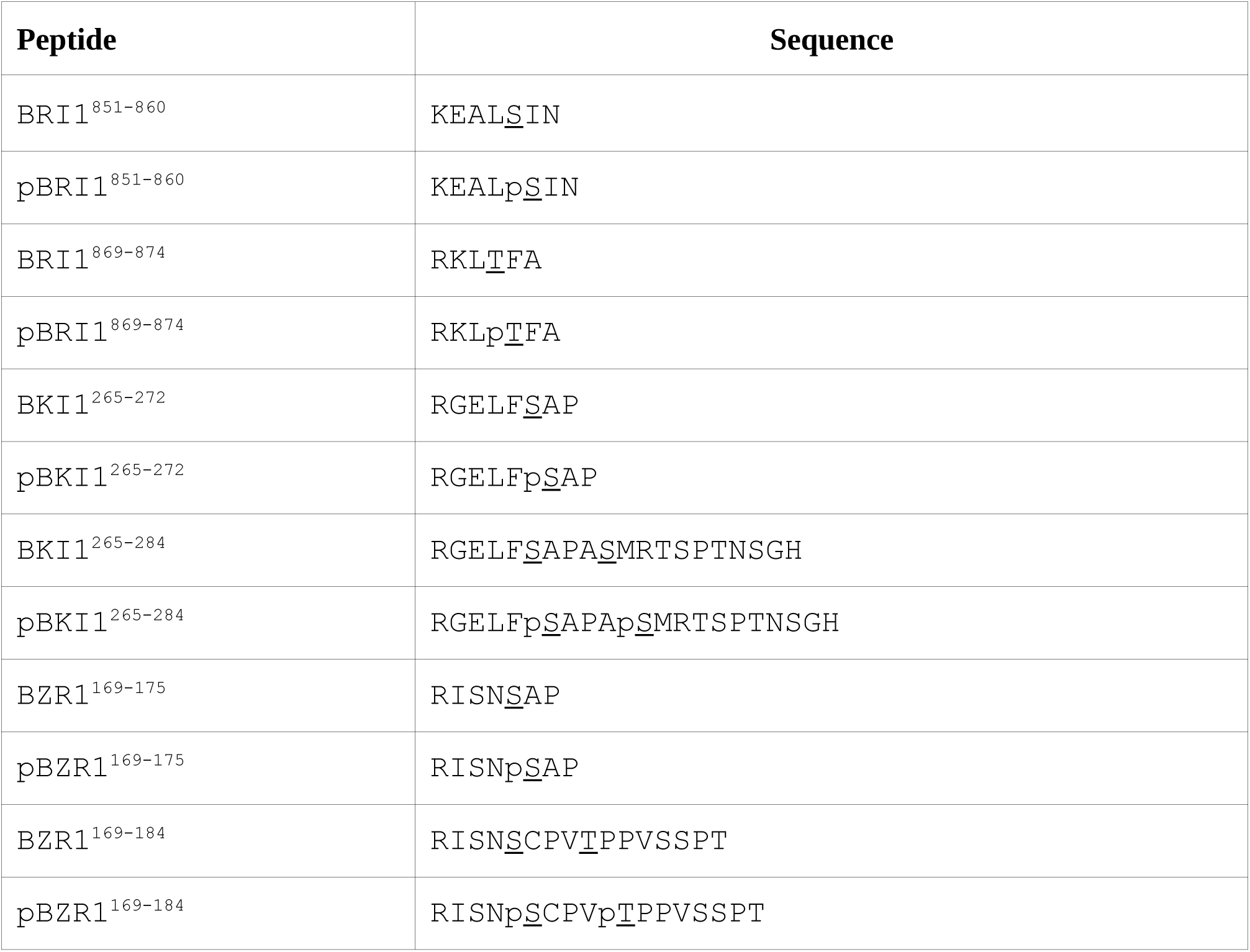
Synthetic peptides used in this work.

Moving downstream of BRI1, the protein kinase inhibitor BKI1 (Wang and Chory, 2006) was previously reported to interact with the 14-3-3κ and 14-3-3 lambda (14-3-3ƛ) isoforms *in planta* and in *in vitro* pull-down assays (Wang et al., 2011). The interaction site was mapped to the C-terminal half of BKI1 (Wang et al., 2011), upstream of the BRI1 docking motif (Jaillais et al., 2011; Wang et al., 2014) and surrounding the phosphorylated Ser270 (Wang et al., 2011). A synthetic phosphopeptide covering Ser270 (RGELF<u>S</u>AP; pBKI1^265-272^) bound to 14-3-3κ with a dissociation constant (K*_D_*) of ∼15 µM and with 2:2 binding stoichiometry (Fig. 1D,E, Table 1). No binding was detected to the unphosphorylated peptide (Fig. 1D,E, Table 1). As phosphorylation of the neighboring Ser274 promotes interaction with 14-3-3s *in vivo* (Wang et al., 2011), we also tested a longer peptide that includes both Ser270 and Ser274 (RGELF<u>S</u>APA<u>S</u>MRTSPTNSGH; BKI1^265-284^). No binding was detected to either double phosphorylated or unphosphorylated versions of this peptide, suggesting that pBKI1^265-272^ represents the minimal 14-3-3 binding motif in BKI1 (Fig. 1D, Table 1).

At the level of BR transcription factors, 14-3-3κ bound a linear motif in BZR1 (RISNp<u>S</u>AP; BZR1^169-175^), which has been previously shown to be phosphorylated by BIN2 (Bai et al., 2007; Gampala et al., 2007), with a dissociation constant of ∼0.5 µM and with 2:2 binding stoichiometry (Fig. 1D,F, Table 1). The motif is located in a potentially unstructured region C-terminal of the BZR1 basic helix-loop-helix DNA binding domain (Nosaki et al., 2018), and upstream of the dominant *bzr1*-*D* missense mutation (Pro234→Leu) (Wang et al., 2002) and the BIN2-docking motif (Peng et al., 2010) (Fig. 1G). In agreement with earlier reports (Bai et al., 2007; Gampala et al., 2007; Tang et al., 2011), 14-3-3κ – BZR1 association was strictly dependent on phosphorylation of Ser173 (Fig. 1D,F). Extension of the BZR1^169-175^ motif to include this second phosphorylation site reduced binding ∼10fold (RISNp<u>S</u>CPVp<u>T</u>PPVSSPT; BZR1^169-184^; Fig. 1D,F, Table 1).

Taken together, three linear phosphopeptide motifs in the receptor kinase BRI1, in the kinase inhibitor BKI1 and in the transcription factor BZR1 bind 14-3-3κ from Arabidopsis with moderate to high affinity.

### BR signaling components show little 14-3-3 isoform preference

Previously identified interactions with BR signaling components in Arabidopsis have been reported for 14-3-3 isoforms from the non-ε group (Fig. 2A). To test if BKI1 or BZR1 show any isoform binding preference, we expressed and purified 14-3-3ƛ and 14-3-3 omega (14-3-3ω) from this sub-group and tested for interaction with the linear motifs defined for BKI1 and BZR1. We found that pBKI1^265-272^ bound 14-3-3ƛ and 14-3-3ω with slightly lower dissociation constants when compared to 14-3-3κ (Fig. 2B). All three isoforms interacted with the core pBZR1^169-175^ motif with highly similar binding constants (Fig. 2C), in agreement with earlier qualitative assays (Wang et al., 2013).

**Fig. 2.**
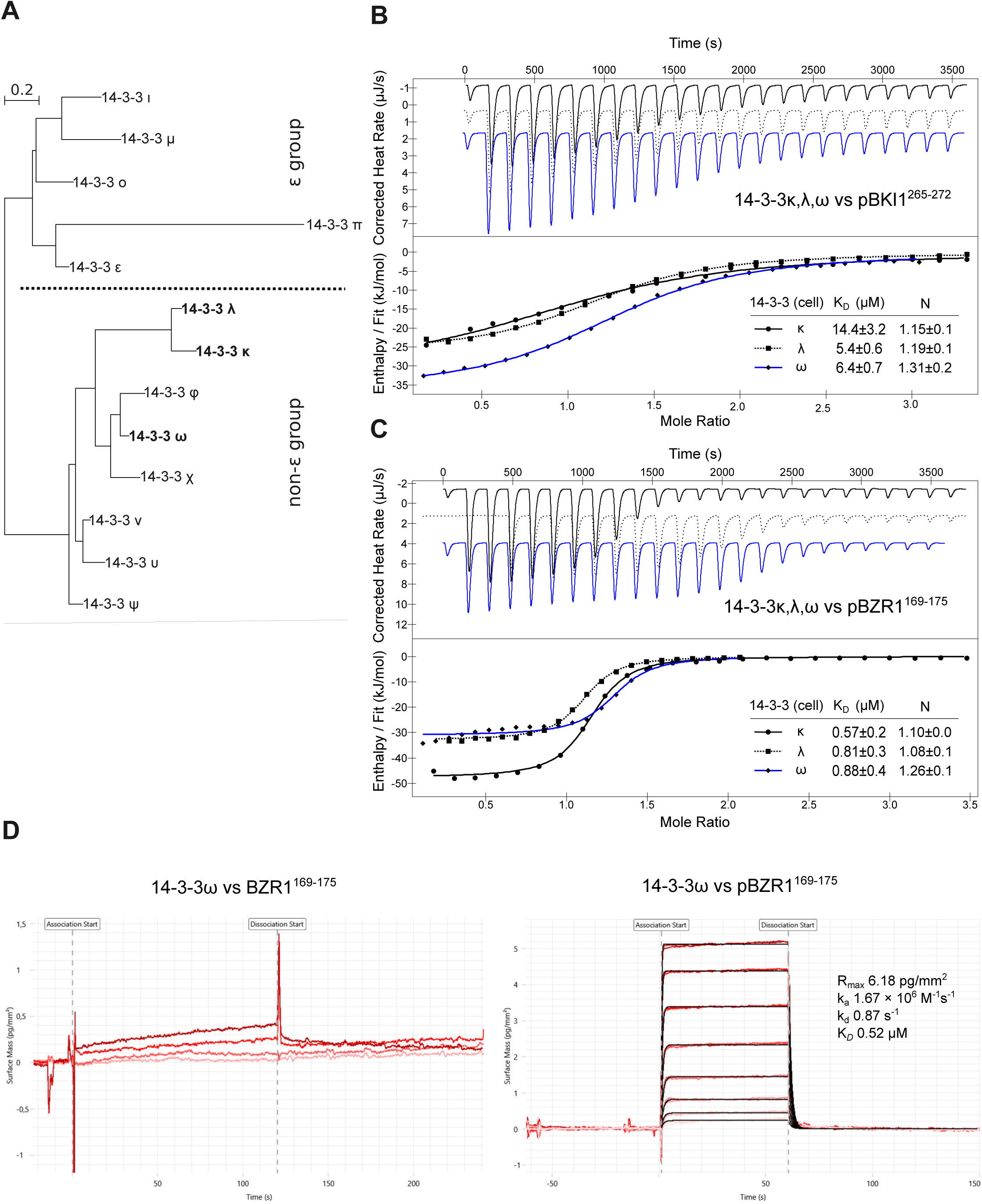
BR signaling components show no 14-3-3 isoform preference. (A) Phylogenetic tree of the 13 14-3-3 isoforms annotated in the Arabidopsis genome. The κ, ƛ and ω isoforms from the non-ε group used for biochemical and crystallographic experiments are highlighted in bold face. A dotted line separates the ε from the non-ε group. The tree was calculated with phyml (Guindon et al., 2010) from a multiple protein sequence alignment of all At14-3-3 isforms generated with MUSCLE (Edgar, 2004) and plotted with the program NjPlot (Perrière and Thioulouse, 1996). (B) Isothermal titration calorimetry of 14-3-3κ (continuous line), 14-3-3ƛ (dotted line) and 14-3-3ω (blue line) vs. the BKI1 minimal binding motif (residues 265-272). Shown are integrated heat peaks (upper panel) vs. time and fitted binding isotherms vs. molar ratio of peptide ligand (lower panel). Table summaries for dissociation constants (K_D_) and binding stoichiometries (N) are shown (±fitting error). (C) Binding of 14-3-3κ, 14-3-3ƛ and 14-3-3ω to the minimal motif in BZR1 (residues 169-175, plotted as in (B)). (D) Grating coupled interferometry (GCI) binding kinetics of 14-3-3 ω vs. BZR^169-175^. Shown are sensorgrams with raw data in red and their respective fits in black. Binding kinetics were analyzed by a 1-to-1 (2:2) binding model. Table summaries of kinetic parameters are shown alongside (ka, association rate constant; kd, dissociation rate constant; K*_D_*, dissociation constant).

To compare our steady-state binding results with a kinetic method, we assayed 14-3-3ω – binding kinetics with grating-coupled interferometry (GCI). We found that the phosphorylated pBZR1^169-175^, but not the un-phosphorylated BZR1^169-175^ peptide bound the surface-adsorbed 14-3-3ω dimer with a dissociation constant of ∼0.5 µM, very similar to the value obtain by ITC (Fig. 2C,D). Binding in GCI is characterized by a relatively fast dissociation rate, with an estimated 14-3-3 – phosphopeptide complex lifetime of only ∼1s (Fig. 2D).

Taken together, the minimal 14-3-3 binding motifs of BKI1 and BZR1 show little isoform preference within the non-ε group in Arabidopsis.

### pBKI1 and pBZR1 are type II 14-3-3 binding motifs

To gain insight into the 14-3-3 binding mechanisms of different BR components, we next performed co-crystallization experiments of full-length 14-3-3κ, 14-3-3ƛ and 14-3-3ω in presence of pBRI1^869-874^, pBKI1^265-272^, pBZR1^169-175^ or the longer pBZR1^169-184^ peptides (see Methods, Table 1). We obtained poorly diffracting crystals for a variety of combinations, and refined structures of 14-3-3ƛ - pBZR1^169-175^ and 14-3-3ω - pBZR1^169-184^ to 2.8 and 3.5 Å resolution, respectively (Table 2). We found the C-terminal 20 amino-acids in 14-3-3ω to be disordered and thus crystallized a truncated 14-3-3ω^1-237^ in complex with pBKI1^265-272^ and pBZR1^169-175^, yielding better crystals diffracting to 2.35 and 1.90 Å resolution, respectively (Table 2). Both 14-3-3ƛ and 14-3-3ω form the canonical 14-3-3 homodimer in the different crystal forms (Fig. 3A). Each protomer is bound to one pBZR1^169-175^ peptide ligand, consistent with the binding stochiometries observed by ITC (Fig. 3A, compare Fig. 1). Dimers within the same asymmetric unit are highly similar, but the ω isoform dimer adopts a more closed conformation when compared to 14-3-3ƛ, even when bound to the same peptide ligand (Fig. 3A). In agreement with our structures, 14-3-3ω forms stable homodimers in solution, as concluded from SEC-RALS (size-exclusion chromatography coupled to right-angle light scattering) experiments (Fig. 3B).

**Fig 3.**
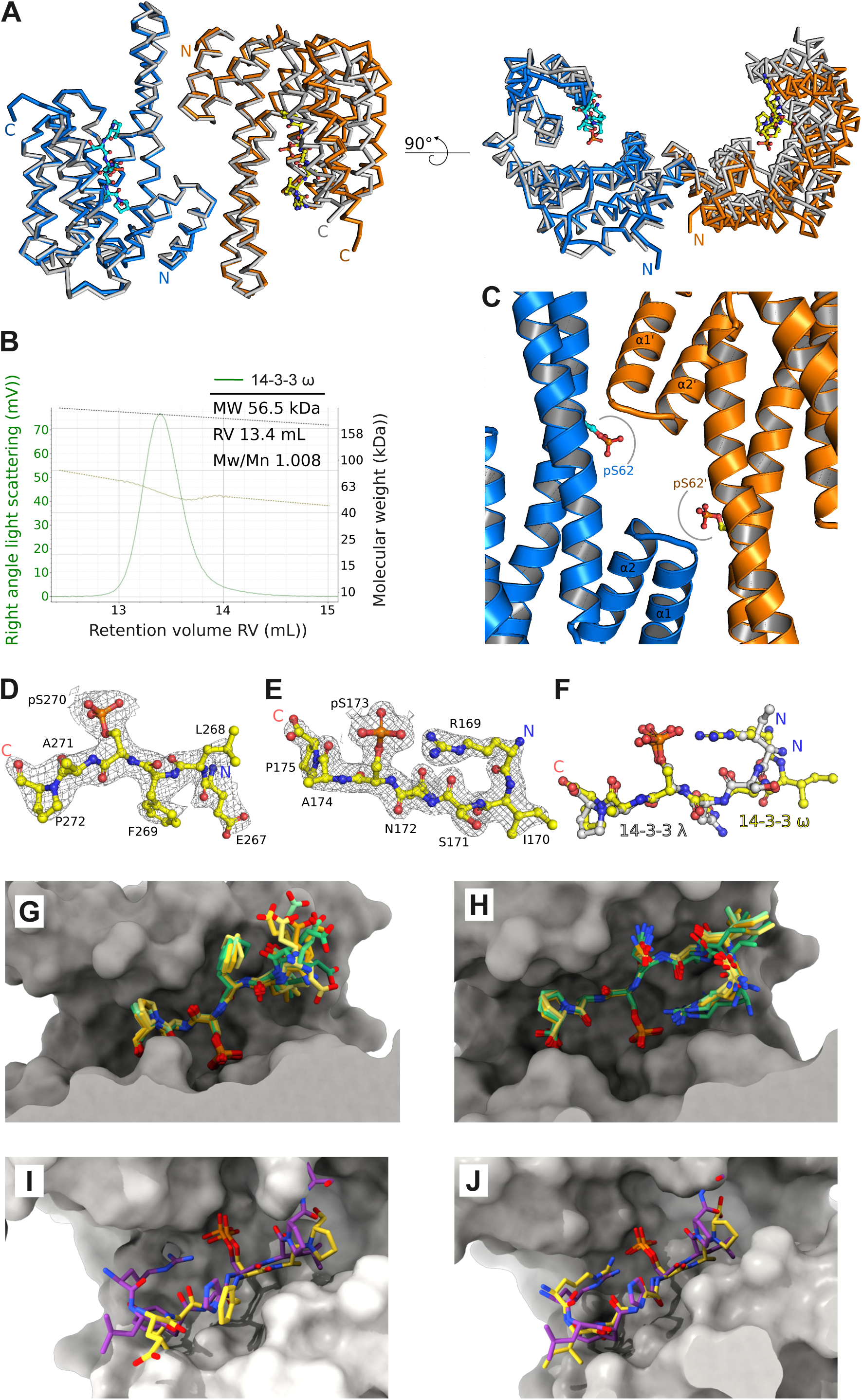
Crystal structures of 14-3-3ƛ and 14-3-3ω reveal type II motif binding modes for pBKI1 and pBZR1. (A) Front and rotated side view of a structural superposition of the 14-3-3ƛ and 14-3-3ω homodimers, each bound to two pBZR1^169-175^ peptides. The two molecules (shown as C_ɑ_ traces) forming the ƛ isoform dimer are colored in blue and orange, respectively, the 14-3-3ω superimposes with an r.m.s.d. (root mean square deviation) of ∼1.6 Å comparing 485 corresponding C_ɑ_ atoms (in gray). The pBZR1 peptides in the ƛ isoform are shown alongside (in bonds representation). (B) Size-exclusion chromatography coupled to right-angle light scattering (SEC-RALS) raw scattering trace of the apo 14-3-3ω isoform (in green) and including the derived molecular masses (light green) of the homodimer. Table summaries report the observed molecular weight (MW), column retention volume (RV) and the dispersity (Mw/Mn). The calculated theoretical molecular weight for At14-3-3ω is ∼58.3 kDa. (C) View of the 14-3-3ω dimer interface (blue and orange ribbon diagrams) containing the previously reported Ser62 (in bonds representation), phosphorylation of which controls dimer-to-monomer transitions in Arabidopsis (Denison et al., 2014). Gray lines indicate potential steric clashes of the phosphorylated amino-acid side chain with the ɑ1-ɑ2 loop in each protomer. (D) Structure of the pBKI1^265-272^ peptide (in yellow, in bonds representation) bound to 14-3-3ƛ with the final (2*F*_o_ – *F*_c_) map contoured at 1.2σ. (E) Structure of pBZR1^169-175^ bound to 14-3-3ω with the final (2*F*_o_ – *F*_c_) map contoured at 1.5σ. (F) Structural superposition of pBZR1^169-175^ bound to 14-3-3ƛ (in gray, in bonds representation) or to 14-3-3ω (in yellow). The 14-3-3ƛ and 14-3-3ω isoform dimers superimpose with a r.m.s.d. of ∼0.7 Å comparing 402 corresponding C_ɑ_ atoms. (G) Structural superposition of all pBKI1^265-272^ peptides bound to the ten 14-3-3ω molecules in the asymmetric unit with a r.m.s.d. of ∼0.5 Å over all atoms. Shown is a molecular surface view of the 14-3-3ω ligand binding site (in gray) with the pBKI1 peptides colored from yellow to green (in bonds representation). (H) Structural superposition of the different pBZR1 peptides in the 14-3-3ω – pBZR1^169-175^ complex (colors as in panel G). (I) Structural superposition of the 14-3-3ω pBKI1^267-272^ (in yellow, in bonds representation) and the Hs14-3-3ζ – type II peptide motif complex (in purple) from a synthetic library (PDB-ID 1qja, r.m.s.d is ∼0.5 Å comparing 195 corresponding C_ɑ_ atoms. (Rittinger et al., 1999). (J) The same comparison as in (I) for the pBZR1^169-172^ peptide.

**Table 2.**
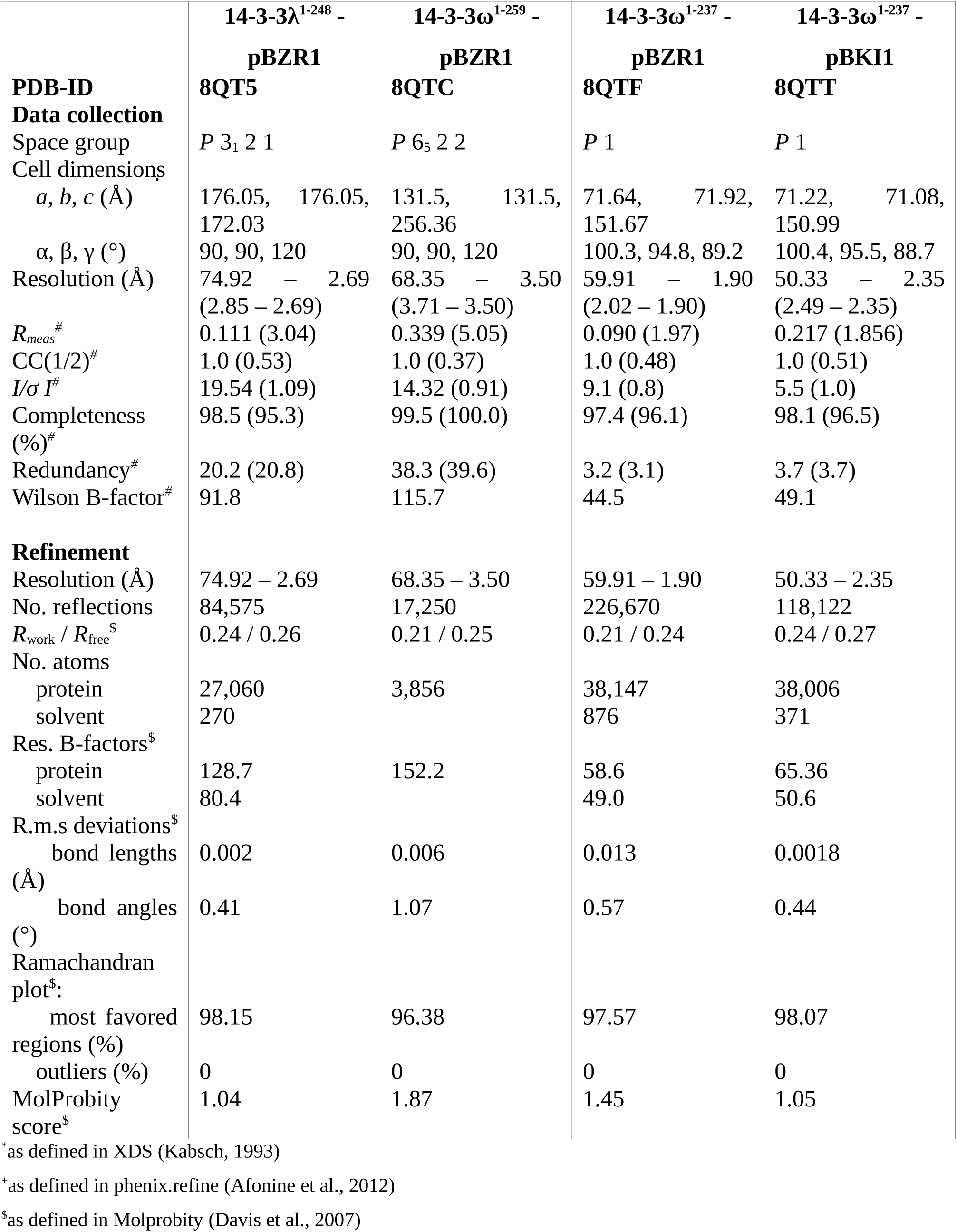
Crystallographic data collection and refinement statistics.

Ser62, phosphorylation of which has been previously reported to induce dimer-to-monomer transition in Arabidopsis 14-3-3ω (Denison et al., 2014; Gökirmak et al., 2015) forms part of the ω homodimer interface in our crystals (Fig. 3C). Ser62 phosphorylation *in silico* induces clashes with residues from the (ɑ1-ɑ2) loop (residues 18-21) that is part of the N-terminal ɑ-helical hairpin in the neighboring molecule, rationalizing why Ser62 phosphorylation induces 14-3-3ω monomerisation (Fig. 3C).

The structure of 14-3-3ƛ in complex with pBKI1^265-272^ (Table 2) revealed the N-terminal Arg^265^ and Gly^266^ residues in the peptide to be largely disordered (full peptide sequence RGELFp<u>S</u>AP, Fig. 3D). In contrast, the entire pBZR1^169-175^ (RISNp<u>S</u>AP) is well defined in the ligand binding site of 14-3-3ω (Fig. 3E), with the N-terminal Arg^169^-Ile^170^ peptide adopting a different conformation in the 14-3-3ƛ – pBZR1^169-175^ complex (Fig. 3F).

We next made use of the high number of molecules in the asymmetric units of our 14-3-3ω crystal form to study the structural plasticity of pBKI1^265-272^ and pBZR1^169-175^ binding. We found the N-terminal Glu^267^-Leu^268^ peptide to bind in different conformations in the ten 14-3-3ω molecules in our structure (Fig. 3G), consistent with the moderate binding affinity for pBKI1^265-272^ in ITC assays (Fig. 1E). In contrast, pBZR1^169-175^ is not only found well-ordered in our 14-3-3ω complex structure (Fig. 3E), but also binds in a highly similar conformation to all 14-3-3ω molecules in the asymmetric unit (Fig. 3H), in good agreement with the high binding affinity observed by ITC (Fig. 1F) and GCI (Fig. 2D).

Three different binding modes have been previously reported for motifs interacting with 14-3-3 proteins (Muslin et al., 1996; Yaffe et al., 1997; Ottmann et al., 2007). Structural comparison of our pBKI1^267-272^ and pBZR1^169-175^ complexes with previous 14-3-3 – ligand complex structures revealed significant similarity to type II 14-3-3 binding motifs, such as found in a Hs14-3-3ζ complex with a synthetic type II peptide with consensus sequence R*X*Φ*X*-p<u>S/T</u>-*X*P (with Φ representing an aromatic or aliphatic residue) (Rittinger et al., 1999), as previously suggested (Gampala et al., 2007; Wang et al., 2011) (Fig. 3I,J).

Taken together, residues 269-272 surrounding the central Ser270 in AtBKI1 and residues 169-175 in AtBZR1 harboring the BIN2 phosphorylation site Ser173 (Gampala et al., 2007; Ryu et al., 2007) represent minimal type II binding motifs for 14-3-3 proteins.

Based on the common binding modes for pBKI1 and pBZR1, we next identified amino-acids interacting with both phosphopeptides in the different complex structures. We found Arg136 and Tyr137 to form hydrogen bond networks with pS270 in pBKI1 and with pS173 in AtBZR1, respectively (Fig. 4A). Asn233 forms hydrogen bonds with the back-bone of both peptides (Fig. 4A). Mutation of Asn233 to Ala in 14-3-3ƛ had little effect on pBKI1^267-272^ binding in ITC assays, but reduced pBZR1^169-175^ and pBRI1^869-874^ interaction by ∼5fold and ∼2fold, respectively (Fig. 4B, compare Figs. 1,2). In contrast, a 14-3-3ƛ Arg136→Leu / Tyr137→ Phe double mutant showed no detectable binding to any of the peptides tested, further highlighting the crucial contribution of threonine/serine phosporylation to AtBRI1, AtBKI1 and AtBZR1 recognition by 14-3-3 proteins (Fig. 4C, compare Fig. 1). The mutant protein has no tendency to aggregate and behaves similar to wild type as a stable homodimer in solution, suggesting that the mutations do not interfere with protein folding or homodimer formation (Fig. 4D).

**Fig. 4.**
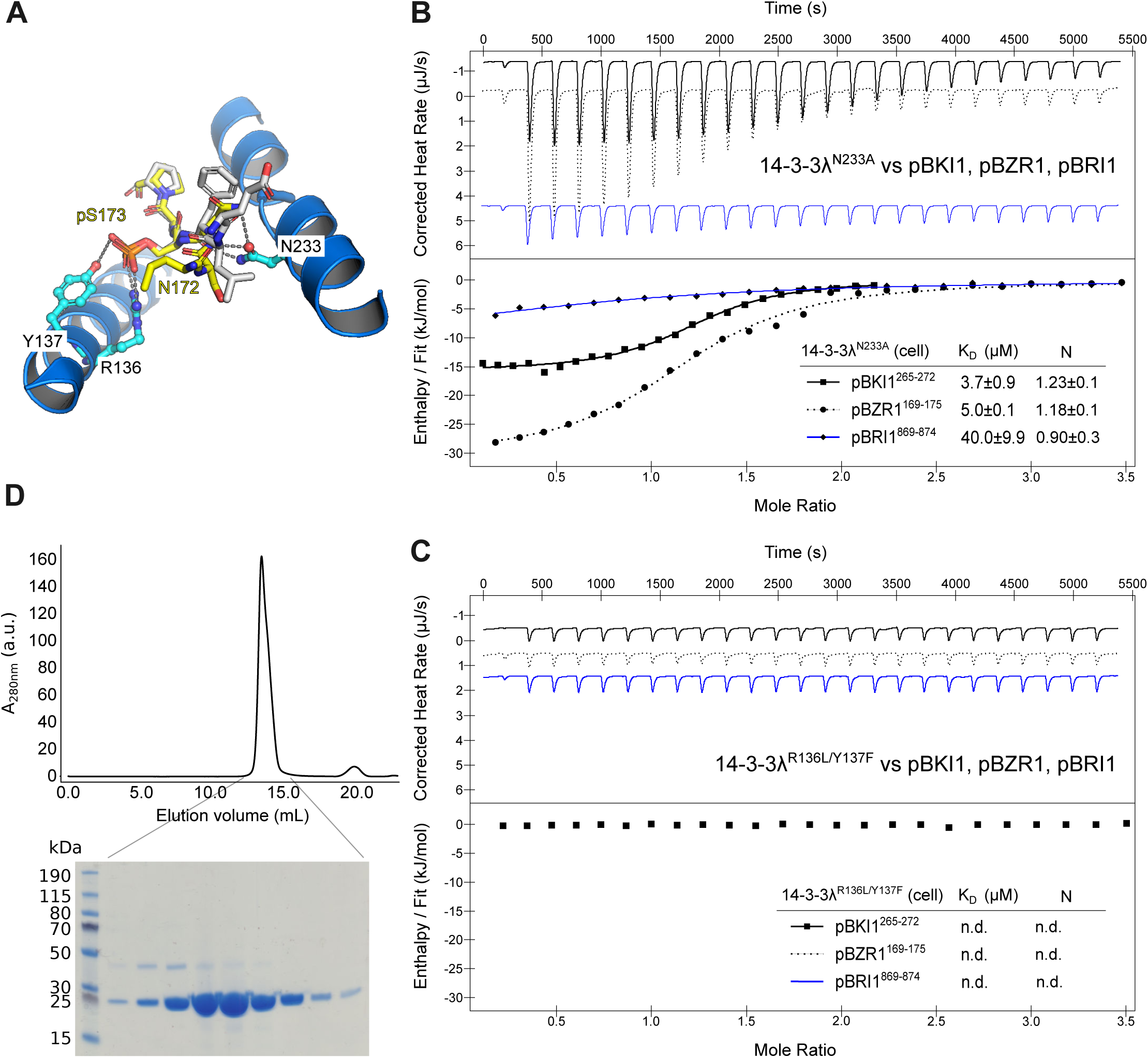
Mutations in the 14-3-3ƛ ligand binding site interfere with pBKI1, pBZR1 and pBRI1 binding. (A) Close up view of pBZR1 (in yellow, in bonds representation) and pBKI1 (in gray) in the 14-3-3 binding groove. The side chains of Arg136, Tyr137 and Asn233 are shown as ball-and-stick models, dashed lines indicate hydrogen bonds (in gray, distance cut-off 3.0Å). (B) Isothermal titration calorimetry of the 14-3-3ƛ Asn233→ Ala mutant vs. pBKI1^265-272^ (continuous line), pBZR1^169-175^ (dashed line) and pBRI1^869-874^ (continuous blue line). Shown are integrated heat peaks (upper panel) vs. time and fitted binding isotherms vs. molar ratio of peptide ligand (lower panel). Table summaries for dissociation constants (K_D_) and binding stoichiometries (N) are shown (±fitting error). (C) ITC analysis of the 14-3-3ƛ Arg136→ Leu / Tyr137 → Phe double mutant. Labels and colors as in panel B. (D) Analytical size exclusion chromatography of the 14-3-3ƛ^R136L/Y137F^ mutant. A Coomassie-stained SDS-PAGE of the homodimeric peak fractions is shown below.

Together, our mutational analysis of 14-3-3ƛ highlights specific binding of the pBRI1^869-874^, pBKI1^267-272^ and pBZR1^169-175^ linear motifs to 14-3-3s from Arabidopsis and validates our pBKI1 and pBZR1 crystal complex structures.

### 14-3-3 non-ε group isoforms are negative regulators of BR signaling

To gain further insight into the contribution of 14-3-3 proteins to BR signaling, we assayed previously generated double and quadruple 14-3-3 isoform loss-of-function mutants (van Kleeff et al., 2014) for BR-related phenotypes. To this end, we used aa established quantitative hypocotyl growth assay in the presence and absence of the brassinosteroid biosynthesis inhibitor brassinazole (BRZ) (Fig. 5) (Asami et al., 2000; Hohmann et al., 2018a, 2018b, 2020). While the κƛ, υν and ɸχ double-mutants behaved similar to wild type, the κƛɸχ, κƛυν and υνɸχ quadruple mutants all displayed gain-of-function phenotypes that were similar to the *bir3-2* control (Fig. 5) The receptor pseudokinase BIR3 is a known negative regulator of BR signaling, keeping the BRI1 receptor from interacting with SERK family co-receptors (Imkampe et al., 2017; Hohmann et al., 2018a). Together, analysis of higher order loss-of-function mutants define 14-3-3 isoforms from the non-ε group as overall negative regulators of BR signaling in Arabidopsis.

**Fig. 5.**
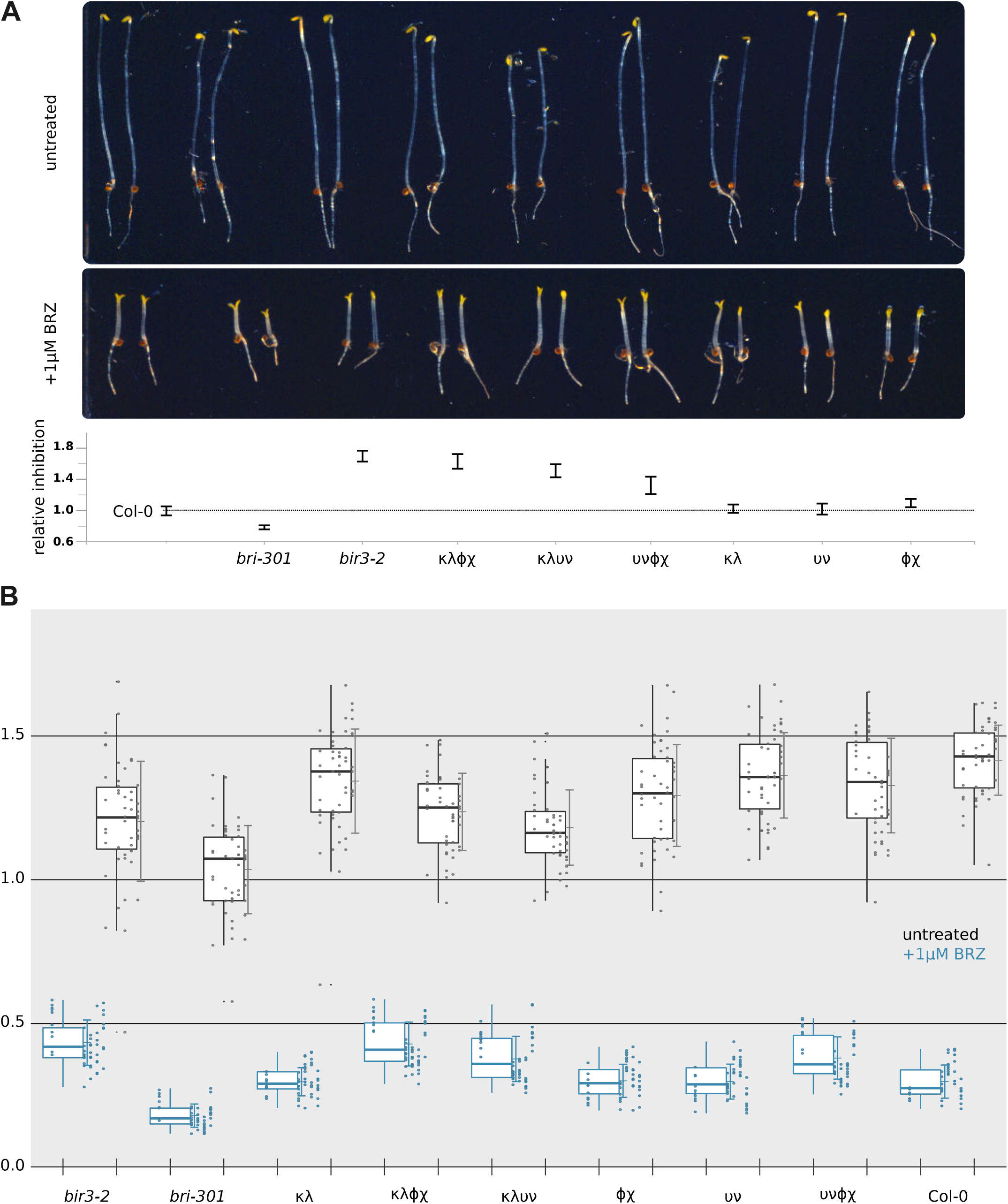
14-3-3 knock-out mutants from the non-ε group show BR gain-of-function phenotypes. (A) Hypocotyl growth assay of dark grown seedlings in the presence and absence of the brassinosteroid biosynthesis inhibitor brassinazole (BRZ). Shown are the growth phenotypes of different non-ε group 14-3-3 loss-of-function double and quadruple mutant combinations compared to the Col-0 wild type, the weak receptor mutant bri1-301 (Xu et al., 2008; Sun et al., 2017; Zhang et al., 2018) and the gain-of-function allele *bir3-2* (Imkampe et al., 2017). Shown below is the quantification of the data with relative inhibition plotted together with lower and upper confidence intervals. For each sample (genotype and treated or untreated) n=50 biologically independent hypocotyls, from five different ½MS plates, were measured. (B) Box plots of the experiment shown in A with raw data depicted as individual dots. Untreated samples shown in black, BRZ treated sample in blue.

## Discussion

Candidate approaches, yeast-2-hybrid screens (Bai et al., 2007; Gampala et al., 2007; Ryu et al., 2007), affinity purifications followed by mass spectrometry (Karlova et al., 2006; Wang et al., 2011), and proximity labeling (Kim et al., 2023) have yielded 14-3-3 interactions with several BR signaling components. In this work, we could confirm and quantify the interaction of 14-3-3s with the receptor BRI1, the inhibitor protein BKI1 and the BR transcription factor BZR1. No interaction of full-length BRI1, BAK1, BSU1 or BIN2 was observed, but it is possible that either critical phosphorylation events are missing in our heterologous expression systems, that interaction with 14-3-3κ is not strong enough to be quantified by ITC, or that interactions sites only become accessible upon interaction with other BR pathway components such as scaffolding proteins (Ehsan et al., 2005; Chaiwanon et al., 2016; Amorim-Silva et al., 2019). It is thus likely that additional interaction sites for 14-3-3 proteins in the BR pathway remain to be identified.

Mapping of the 14-3-3 binding sites yielded a new motif in the N-terminal lobe of the BRI1 kinase domain surrounding Thr872 (Fig. 1B,C). Thr872 represents a BRI1 (trans-) autophosphorylation site (Oh et al., 2000; Wang et al., 2005). Mutation of Thr872 → Ala increases the catalytic activity of BRI1 *in vitro* and has a growth-promoting effect *in planta*. Based on our observation that 14-3-3s can only bind BRI1^869-874^ when phosphorylated at Thr872, we speculate that the gain-of-function effect of the BRI1 Thr872→ Ala mutant could in part be explained by the lack of interaction between 14-3-3 proteins and the mutant BRI1 kinase domain (Wang et al., 2005, 2008). Since reciprocal BRI1 – SERK transphoshorylation appears to be driven by spacial proximity, 14-3-3s may also sterically hinder receptor – co-receptor interaction in the cytoplasm, thereby negatively regulating BR signaling at the level of the receptor complex (Wang et al., 2008; Santiago et al., 2013; Bojar et al., 2014; Hohmann et al., 2018a, 2020).

Biochemical mapping of the 14-3-3 binding site in BKI1 yielded a linear motif that is somewhat shorter than previously envisioned, yet contains Ser270, which is likely phosphorylated by BRI1 to release BKI1 from the plasma membrane (Wang et al., 2011) (Fig. 1D,E). It is of note that a longer peptide additionally containing BKI1 Ser274 does not bind to 14-3-3κ *in vitro* (Fig 1. D). However phosphorylation of both Ser270 and Ser274 are required for BKI1 membrane dissociation (Wang et al., 2011). It is thus possible that 14-3-3-dependent and -independent mechanisms regulate BKI1’s function, as previously reported (Jaillais et al., 2011; Wang et al., 2011).

We observed tight interaction of BZR1 with different 14-3-3 isoforms involving a linear motif that was previously reported to center around Ser173, a BIN2 phosphorylation site (Wang et al., 2002; Yin et al., 2002; Vert and Chory, 2006; Gampala et al., 2007). Again, BZR1^169-175^ binding to 14-3-3s is strictly phosphorylation dependent, rationalizing why protein phosphatase 2A-mediated BZR1 dephosphorylation abolishes 14-3-3 binding (Tang et al., 2011) (Fig. 1F). The adjacent Thr177 represents a putative BZR1 phosphorylation site that affects 14-3-3 interaction (Wang et al., 2002; Ryu et al., 2010). Importantly, inclusion of this site in a longer BZR1 peptide strongly reduces binding, suggesting that differential BZR1 phosphorylation may both promote and inhibit interaction with 14-3-3 proteins (Fig. 1F). Alternatively, recently identified scaffolding proteins may enable more complex interaction networks *in vivo* (Li et al., 2023). The position of the defined 14-3-3 binding motif in BZR1 in relation to the DNA binding motifs however makes it difficult to deduce the 14-3-3 regulatory mechanism in BZR1 (Fig. 1G). It remains possible that 14-3-3 mediated cytosolic retention of BZR1 represents the central regulatory event *in planta* (Gampala et al., 2007; Lozano-Durán et al., 2014; Ryu et al., 2007; Tang et al., 2011; Yu et al., 2023). The range of binding affinities (∼0.5 - ∼20 µM) observed for the various 14-3-3 – BR component interactions is comparable to what has been previously reported for some (Gao et al., 2014), but not all (Fuglsang et al., 2003; Latz et al., 2007; Ottmann et al., 2007) 14-3-3 – protein interactions in plants.

Our biophysical and crystallographic studies define the minimal binding motifs for BKI1 and BZR1, reveal both fragments to represent typical type II motifs (Rittinger et al., 1999) and uncover conformational changes in Arabidopsis 14-3-3 homodimers (Yang et al., 2006). Similar conformational transitions have been previously observed for human 14-3-3 isoforms (Benzinger et al., 2005; Yang et al., 2006), where they may contribute to ligand binding specificity (Modi et al., 2020). Ligand specificity may also be regulated by 14-3-3 monomerisation (Fig. 3C) (Denison et al., 2014; Gökirmak et al., 2015), or by heteromer formation between different 14-3-3 isoforms, but it is presently unknown to what extend these assemblies exist *in planta*.

Quantitative hypocotyl growth assays suggest an overall function for 14-3-3 proteins from the non-ε group as negative regulators of BR signaling, potentially highlighting a key function for 14-3-3 proteins in BZR1 nucleo-cytoplasmic partitioning (Fig. 5). Different phenotypes have been previously reported for 14-3-3 isoforms from the ε group, suggesting that a functional differentiation of different 14-3-3 isoforms within the BR pathway may yet exist (Lee et al., 2020). The gain-of-function phenotypes observed in the 14-3-3 κƛɸχ, κƛυν quadruple mutants (van Kleeff et al., 2014) are similar to *bir3* mutants (Imkampe et al., 2017; Hohmann et al., 2018a), indicating that 14-3-3 proteins may be important but not essential components of BR signaling. We envision that 14-3-3 proteins regulate BR signaling by several mechanisms including regulation of enzyme activity or substrate binding at the level of the receptor complex, other protein – protein interactions and protein sub-cellular localization in the case of BKI1 and BZR1.

## Materials and Methods

### Protein expression and purification

Full-length At14-3-3-κ^1-248^ (Uniprot-ID P48348, http://uniprot.org), At14-3-3-ƛ^1-248^ (P48349), At14-3-3-ω^1-259^ (Q01525) or C-terminal truncated At14-3-3-κ^1-240^, At14-3-3-ƛ^1-240^,or At14-3-3-ω^1-237^ isoforms and BIN2^7-380^ from Arabidopsis were cloned from synthetic genes codon-optimized for expression in *E. coli* (Geneart, Thermofisher) into vector pMH-HStrxT providing an N-terminal thioredoxin A (trxA) fusion protein containing 8xHis and StrepII affinity tags, and a tobacco etch virus protease (TEV) recognition site. Protein expression in *E. coli* BL21 (DE3) RIL grown to OD_600nm_= 0.6 was induced with 0.2 mM ispropyl β-D-galactoside in terrific broth at 16 °C for 16 h. Cells were collected by centrifugation at 4,500 × g for 30 min, washed in PBS buffer, centrifuged again at 4,500 × g for 15 min and snap-frozen in liquid N_2_. For protein purification cells were resuspended in buffer A (20 mM Tris-HCl pH 8.0, 500 mM NaCl), and lysed by sonication (Branson DS450). The lysate was cleared by centrifugation at 7,000 × g for 60 min and filtrated using a 0.45 µm molecular weight cut-off filter (Pall Corporation). The supernatant was loaded onto a Ni^2+^ affinity column (HisTrap HP 5ml, Cytiva Life Sciences), washed with buffer A and eluted in buffer A supplemented with 250 mM imidazole pH 8.0. The elution fractions containing the trxA-14-3-3 or trxA-BIN2 fusion proteins were loaded onto a 5 ml Strep-Tactin Superflow High Capacity column (IBA Lifesciences), washed with buffer A and eluted in buffer A supplemented with 2.5 mM desthiobiotin. The elution fractions were incubated with TEV for 16 h at 4 °C during dialysis against buffer A. The 8xHis-tagged trxA fusion tag was removed by a second Ni^2+^ affinity step, and the cleaved protein was further purified by gel filtration on a Superdex 75 HR26/60 column (Cytiva Life Sciences), equilibrated in 20 mM Hepes pH 7.5, 150 mM NaCl, 5 mM β-mercaptoethanol. Dimeric (14-3-3s) or monomeric (BIN2) peak fractions were concentrated to 10-40 mg/ml and used directly for biochemistry and crystallization experiments. Point mutations were introduced by site-directed mutagenesis and variants At14-3-3-ƛ^N233A^ and 14-3-3 ƛ^R136L/Y137F^ were purified using the same protocol as described for the wild type.

The BRI1^814-1196^ and BAK1^250-615^/SERK3 cytoplasmic domains were purified as previously described (Bojar et al., 2014). BSK1 (residues 55-512), BSU1 (residues 1-793) and BIN2 (residues 7-380) were cloned into a modified pACEBac1 plasmid (Bieniossek et al., 2012) providing TEV- cleavable N-terminal 8xHis and tandem StrepII affinity tags, and expressed in baculovirus-infected *Trichoplusia ni* Tnao cells (Hashimoto et al., 2010). Cells grown to a density of 1.5 × 10^6^ cells ml^-1^ were infected with 10 ml of virus subjected to two rounds of viral amplification per 250 ml of cells and incubated for 48 h at 28 °C. Cell pellets were harvested by centrifugation at 2,000 × g for 20 min, resuspended in lysis buffer (50 mM Tris pH 8.0, 500 mM NaCl, 2 mM MgCl_2_, 2 mM β-Mercaptoethanol), lysed by sonication (Branson DS450), centrifuged at 60,000 x g for 45 min and loaded on a Ni^2+^ affinity column (Ni Sepharose Excel 5ml, Cytiva Life Sciences). The column was washed with 10 column volumes of lysis buffer and 8xHis tagged target proteins were eluted in lysis buffer supplemented with 500 mM Imidazole pH 8.0. Elution fractions were dialyzed against Strep buffer (50 mM Tris pH 8.0, 150 mM NaCl, 1 mM EDTA) and loaded onto a 5 ml Strep-Tactin Superflow High Capacity colum (IBA Lifesciences). Proteins were eluted from the column in Strep buffer supplement with 2.5 mM desthiobiotin followed by over-night TEV cleavage of the N-terminal affinity tag for 16 h at 4 °C during dialysis against Strep buffer. Cleaved proteins were separated from the tandem affinity tag by a second Strep affinity chromatography step and directly used for biochemical assays.

### Isothermal titration calorimetry (ITC)

All ITC experiments were performed on a Nano ITC (TA Instruments) with a 1.0 ml standard cell and a 250 μl titration syringe at 25 °C. Proteins were gelfiltrated or dialyzed into ITC buffer (25 mM Hepes pH 7.0, 50 mM KCl) prior to all experiments. Synthesized peptide ligands (Peptide Specialty Laboratories, Heidelberg, Germany) were dissolved in the same buffer. 10 μl of the respective peptide or phosphopeptide ligand (Table 1) at a concentration of 600 μM was injected into 50 μM 14-3-3 protein solution in the cell at 150 s intervals (25 injections). Data was corrected for the dilution heat and analyzed using NanoAnalyze® program (version 3.5) as provided by the manufacturer. All ITC assays were performed at least twice.

### Grating Coupling Interferometry

The binding kinetics of 14-3-3ω vs the BZR1^169-175^ and pBZR1^169-175^ peptides (Table 1) were assessed on a Creoptix® WAVE system using a streptavidin-coated 4PCH WAVEchip® (long polycarboxylate matrix; Creoptix AG, Switzerland). Chips were first conditioned with 100 mM sodium borate pH 9.0, 1 M NaCl. Next, after the activation of the chip surface with a 1:1 mix of 400 mM N-(3-dimethylaminopropyl)-N’-ethylcarbodiimide hydrochloride and 100 mM N-272 hydroxysuccinimide (Xantec, Germany), streptavidin was immobilized on the chip surface through the injection of 50 µg/ml streptavidin (Thermo Fisher Scientific 43-4301) in 10 mM sodium acetate pH 5.0 until high density (∼ 10,000 pg/mm^2^) was reached. Biotinylation of 14-3-3ω for capturing onto streptavidin chip was performed by mixing equimolar amounts of protein and biotin (EZ-Link™ NHS-Biotin ThermoFisher Scientific). 170 µM of 14-3-3ω were incubated on ice with 170 µM of biotin (previously dissolved in water to 20 mM) for 1.5 min. The biotinylation was performed in 100 mM Hepes pH 8 and 150 mM NaCl. The biotinylated 14-3-3ω dimer was purified by size exclusion chromatography in 20 mM Hepes pH 7.5 and 150 mM NaCl on a Superdex 10/300 increase column (Cytiva Life Sciences) to verify the homogeneity and to remove excess biotin. Gel filtrated biotinylated 14-3-3ω (0.45 mg/ml) was directly immobilized on the streptavidin-coated chip until a density of about 500 pg/ mm^2^ was reached. The synthetic BRZ1^169-175^ and pBZR1^169-175^ peptides were dissolved in 20 mM Hepes and 150 mM NaCl buffer to a final concentration of 10 mM. Kinetic analysis of 14-3-3ω and BZR1^169-174^ / pBZR1^169-174^ interactions was performed at 25°C by flushing a dilution series of the analyte pBZR1 with a 2 dilution factor starting from 10 µM. Data analysis and data fitting were done using the Creoptix WAVEcontrol® software.

### Analytical size-exclusion chromatography

Gel filtration experiments were performed using a Superdex 200 increase 10/300 GL column (GE Healthcare) pre-equilibrated in 25 mM Hepes pH 7.0, 50 mM KCl. 500 μl of the respective protein (2.0 mg/ml) was loaded sequentially onto the column, and elution at 0.75 ml/ml was monitored by ultraviolet absorbance at 280 nm. Peak fractions were analyzed by SDS–PAGE gel electrophoresis followed by Coomassie staining.

### Right-angle light scattering

The oligomeric state of the 14-3-3ω isoform was analyzed by size exclusion chromatography coupled to right angle light scattering (SEC-RALS), using an OMNISEC RESOLVE / REVEAL combined system (Malvern Panalytical). Instrument calibration was performed with a BSA standard (Thermo Scientific Albumin Standard). 20 μM 14-3-3ω in a volume of 50 μl was separated on a Superdex 200 increase 10/300 GL column (Cytiva Life Sciences) in 25 mM Hepes pH 7.0, 50 mM KCl, at a column temperature of 35 °C and a flow rate of 0.7 ml min^-1^. Data were analyzed using the OMNISEC software (version 10.41).

### Protein crystallization

Hexagonal crystal of 14-3-3λ developed at room temperature from hanging drops composed of 1.5 μL of protein solution (14-3-3λ at 32 mg/ml in the presence of 1 mM pBZR ^169-175^, Table 1) and 1.5 μL of crystallization buffer (22 % [w/v] PEG 10,000, 0.1 M ammonium acetate, 0.1 M Bis-Tris [pH 5.5]) suspended over 1.0 ml of the latter as reservoir solution. Crystals were improved in several rounds of micro-seeding and then transferred in reservoir solution supplemented with 20% (v/v) glycerol and snap-frozen in liquid N_2_. Crystals of 14-3-3ω^1-259^ (35 mg/ml and in the presence of 3 mM pBZR1^169-184^, Table 1) developed in 2 M (NH_4_)_2_SO4, 10 mM CoCl_2_ · 6 H_2_O, 0.1 M Mes (pH 6.5) and were snap-frozen in crystallization buffer containing a final concentration of 20 % (v/v) glycerol. Triclinic crystals of 14-3-3ω^1-237^ (25 mg/ml) and in the presence of either 3 mM pBZR1^169-175^ or 3 mM pBKI1^265-272^ (Table 1) developed in Morpheus (Molecular Dimensions) condition G8 (with a final precipitant stock concentration of 50% [v/v]) using micro-seeding protocols. Crystals were directly frozen in liquid N_2_. Data processing and scaling was done with XDS (version January, 2022) (Kabsch, 1993).

### Structure solution and refinement

The structures of the different 14-3-3 – peptide complexes were solved by molecular replacement as implemented in the program Phaser (McCoy et al., 2007), using Protein Data Bank (http://rcsb.org) ID 2o98 (Ottmann et al., 2007) as the initial search model. Structures were completed in iterative rounds of manual model-building in COOT (Emsley and Cowtan, 2004) and restrained NCS (non-crystallographic symmetry) refinement in phenix.refine (Adams et al., 2010). Structural superpositions were done using phenix.superpose_pdbs. Inspection of the final models with phenix.molprobity (Davis et al., 2007) revealed excellent stereochemistry (Table 2). Structural representations were done with Pymol (https://sourceforge.net/projects/pymol/) and ChimeraX (Goddard et al., 2018).

### Hypocotyl Growth Assay

Wild-type and 14-3-3 mutant seeds (van Kleeff et al., 2014) were surface sterilized, stratified at 4°C for 2 d, and plated on half-strength Murashige and Skoog (½MS) medium containing 0.8% (w/v) agar and supplemented with 1 μM BRZ from a 10 mM stock solution in 100% DMSO (Tokyo Chemical Industry) or, for the controls, with 0.1% (v/v) DMSO. The *bir3-2* (SALK_116632)3 T-DNA insertion line was obtained from the Nottingham Arabidopsis Stock Center (NASC), the bri1-301 mutant has been described previously (Xu et al., 2008). Following a 1 h light exposure to induce germination, plates were wrapped in aluminum foil and incubated in the dark at 22°C for 5 d. We then scanned the plates at 600 dots per inch resolution on a regular flatbed scanner (CanoScan 9000F; Canon), measured hypocotyl lengths using Fiji (Schindelin et al., 2012), and analyzed the results in R version 4.1 (R Core Team, 2014) using the packages mratios (Kitsche and Hothorn, 2014) and multcomp (Hothorn et al., 2008). Rather than P-values, we report unadjusted 95% confidence limits for fold-changes. We used a mixed-effects model for the ratio of a given line to the wild-type Col-0, allowing for heterogeneous variances, to analyze log-transformed end point hypocotyl lengths. To evaluate treatment-by-mutant interactions, we calculated the 95% two-sided confidence intervals for the relative inhibition (Col-0: untreated versus BRZ-treated hypocotyl length)/(any genotype: untreated versus BRZ-treated hypocotyl length) for the log-transformed length.

## Data availability

Crystallographic coordinates and associated structure factors have been deposited with the Protein Data Bank (http://rcsb.org) with accession numbers 8qt5 (14-3-3λ^1-248^ – pBZR1), 8qtc (14-3-3ω^1-237^ – pBZR1), 8qtf (14-3-3ω^1-237^ – pBZR1) and 8qtt (14-3-3ω^1-237^ – pBKI1).

## Acknowledgments

We thank Jacobo Martinez for his help in the early stages of this project, Bert de Boer (Amsterdam) for providing us with the Arabidopsis 14-3-3 κƛ, υν, ɸχ, κƛɸχ, κƛυν and υνɸχ transgenic lines and Philippe Rieu for carefully reading the manuscript.

## Author Contributions

E.O., U.H. and M.H. designed research. E.O. expressed and purified 14-3-3 proteins, BRI1, BAK1 and BIN2, performed ITC assays and crystallized proteins. A.M. purified BSU1, BIN2 and BSK1 and performed trans-phosphorylation and GCI assays. U.H. and E. O. performed and analyzed hypocotyl growth assays. M.H and E. O. collected X-ray diffraction data and build and refined the crystallographic structures. M.H. wrote the manuscript with input from all authors.

